# Massively parallel reporter assay reveals promoter-, position-, and strand-specific effects in transcription start sites

**DOI:** 10.1101/2025.10.13.659964

**Authors:** Maya Bose, Adelaide Tovar, Yasuhiro Kyono, Arushi Varshney, Jacob O. Kitzman, Stephen C.J. Parker

**Affiliations:** Department of Computational Medicine and Bioinformatics, University of Michigan, Ann Arbor, MI 48109, USA; Department of Human Genetics, University of Michigan, Ann Arbor, MI 48109, USA; Department of Biostatistics, University of Michigan, Ann Arbor, MI 48109, USA

## Abstract

Massively parallel reporter assays (MPRA) are a high-throughput method of assessing the activity of candidate cis-regulatory sequences, and can be used to detect allelic differences at disease-associated variants. Previous MPRA studies have screened thousands of functional SNPs associated with various complex traits and conditions. Most MPRA libraries utilize a single plasmid configuration, a single minimal promoter to drive expression, and a single-strand orientation, which may fail to capture the context-dependent activity of disease-associated cis-regulatory elements. We interrogate the potential regulatory differences introduced by variable MPRA plasmid promoters and positions. We used an MPRA library to quantify the activity of 1,305 pancreatic islet-derived transcription start sites generated from CAP analysis of gene expression profiling. We cloned fragments upstream or downstream of a reporter gene along with either the human insulin (*INS*) promoter or a synthetic housekeeping promoter (SCP1). We used elastic net regression to predict position-specific fragment activity based on enrichment of transcription factor binding site motifs, and generalized linear models to predict position-specific fragment activity from tissue-specific chromatin state regulatory annotations. Our results support the use of MPRA strategies that account for context-dependent factors when assaying candidate regulatory elements in pursuit of understanding complex genetic diseases.

## INTRODUCTION

The controlled expression of individual genes in response to spatio-temporal factors is crucial to proper function of biological processes in living organisms. These programs of gene activation and inactivation are maintained via the non-coding genome, which makes up approximately 98.5% of the human genome and, rather than directly coding for genes, encompasses a wide variety of cis-regulatory elements (CREs) such as enhancers, promoters, and silencers(1). While several of these elements act ubiquitously across cell types to control fundamental biological processes, the ability of many of these elements to act with cell-type specificity is essential to their function. For example, promoters controlling the expression of genes associated with essential cell functions, such as autophagy or DNA replication, need to be active at some point in all cells, regardless of type. Meanwhile, the human insulin (*INS*) promoter controls the expression of the human insulin gene, which is exclusively expressed in the beta cells found in pancreatic islets as a central part of their function (2). Beta cell dysfunction and, in turn, inadequate expression of insulin is associated with several metabolic diseases, including type 2 diabetes mellitus (T2D)(3, 4). Given that more than 90% of T2D GWAS variants are located in non-coding regions of the genome, the most likely mechanism of disease lies in altered gene regulation (5, 6). A more thorough understanding of gene regulation is thus essential to the study of complex disease.

Massively parallel reporter assays (MPRAs) have emerged as a popular tool for measuring the regulatory activity of many sequences in a high-throughput manner(7). Briefly, a library of sequence fragments derived from GWAS peaks or sequencing data are cloned into individual plasmids, each paired with a unique barcode. These plasmids are then transfected into cells, and the regulatory activity of each fragment can be determined by sequencing these cells for DNA and RNA, with activity calculated as the ratio between the two values. Previously, MPRAs have been used to perform saturation mutagenesis in promoters(8) and enhancers(9), decode regulatory grammars(10), predict enhancer activity(11–13), and detect the effects of disease-associated SNPs in regulatory elements(14–16). MPRA experiments are intentionally synthetic in that they test the regulatory activity of sequences outside of their native genomic context. This makes MPRAs a powerful tool for testing the effects of specific variants on gene regulation while controlling for other genomic environmental factors such as interactions with promoters and relative positioning to other CREs.

The plasmid configurations used in MPRA studies may introduce limitations to the interpretability of results. In a review of 118 published MPRA datasets (from 2012 to January 31, 2024; Figure 1A), 91% test fragments alongside a single minimal core promoter. However, given the potential sequence-specificity of some enhancer-promoter links, these assays may mischaracterize enhancers that are active in the cell type being tested, but do not strongly target the core promoter being used in the assay (17, 18). Additionally, 94% of reviewed MPRA experiments test their fragments in a single position relative to their promoter element and sequence orientation (Figure 1B), which likewise may not reflect the full spectrum of the function of these sequences in their native context, especially in the case of testing strongly unidirectional elements such as promoters (19, 20). For example, several studies screening regulatory elements (promoter and enhancers) and their variants for activity only test a single configuration, and thus may be finding false negatives that would be expressed in the alternate configuration (21–23). Currently, few studies test how modular variations in promoter, position, and strandedness of MPRA plasmid configurations may affect readout and thus bias the interpretation of downstream results in the context of cell type and disease (24).

**Figure 1.**
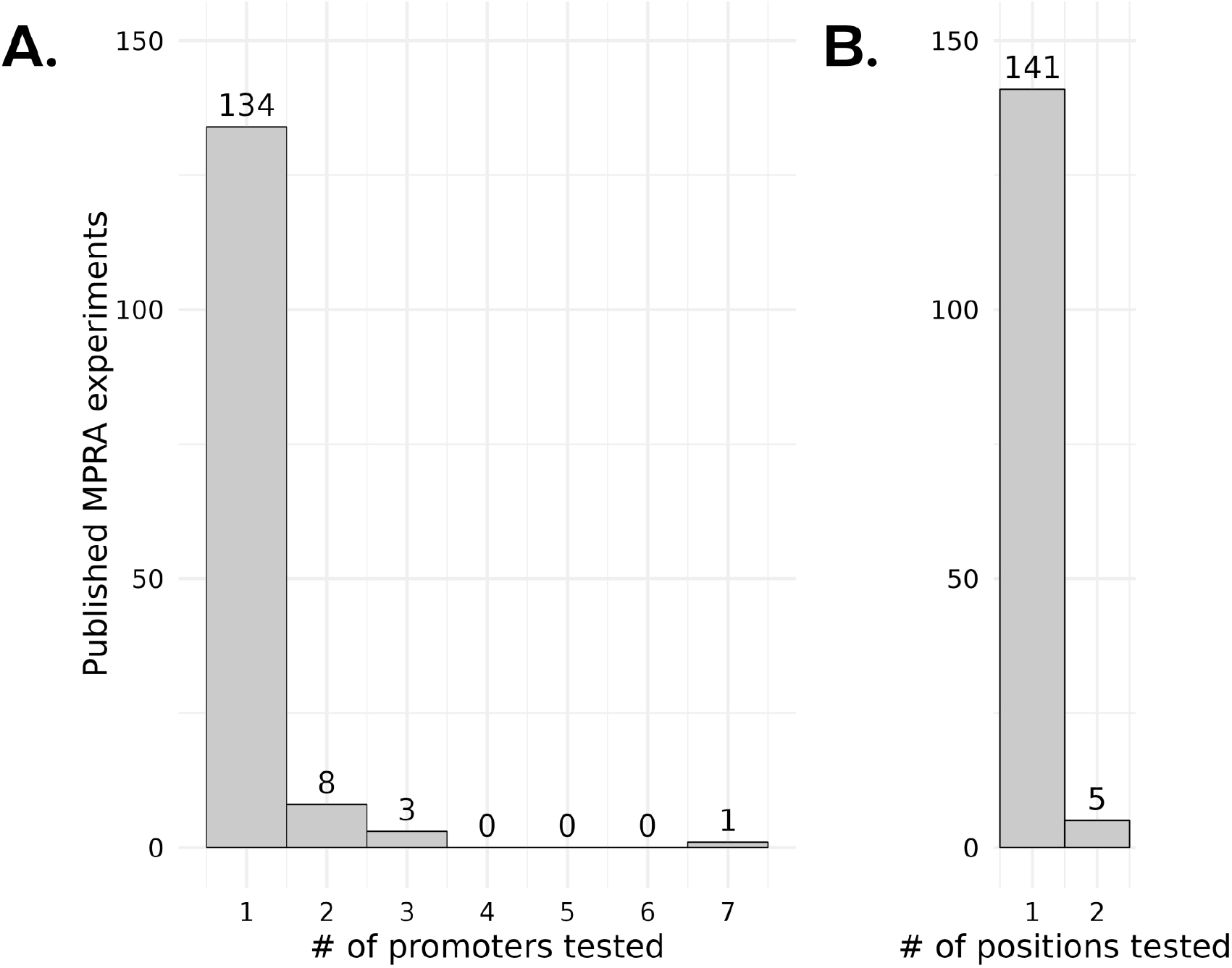
Review of 146 published MPRA experiments (2012 to August 29, 2025). (A) Number of promoters used within a study across all reviewed studies. (B) Number of fragment positions tested within a study across all reviewed studies. A list of these publications is provided in Supplementary Data 1.

In this study, we systematically test 1,305 short fragments (198bp) derived from islet transcription start sites in four different plasmid configurations: using either the super core promoter 1 (SCP1) or the *INS* promoter, and positionally placed either upstream or downstream of said promoter. This set of plasmid configurations was also used to test a set of fragments with T2D GWAS-derived variants in a separate study(24). SCP1 is a synthetic housekeeping promoter composed of multiple core promoter sequences, and is capable of directing RNA polymerase II transcription ubiquitously across cell types(25). As mentioned, the human insulin promoter controls the expression of the human insulin gene (*INS*), and its activity is specific to islets. We transfected our library into the rat insulinoma 823/13 cell line–a model cell line for pancreatic beta cells. We hypothesized that the *INS* promoter and upstream positional configuration, when placed in a disease-relevant cell type, would more accurately reflect the true biological context of the fragments being tested, and would thus exhibit better compatibility and enhanced regulatory activity. While we observed clear promoter- and position-dependent effects across our fragment library, with a clear activity bias for fragments in the upstream position, we also observed a stronger bias for fragments with the SCP1 promoter over the *INS* promoter. We utilized an elastic net regression to identify sets of TF motifs that exhibit significant configuration bias. Finally, we use a generalized linear model to demonstrate the tissue-dependent and strand-specific enhancer activity of our fragments. Cumulatively, the results of these two studies should prompt close consideration of context-specific effects when designing MPRA libraries.

## MATERIAL AND METHODS

### MPRA library construction and cell transfection

Tag clusters were identified from CAGE sequencing libraries from 71 human pancreatic islet samples as described in Varshney et al. (2021). MPRA libraries testing the expression of fragments derived from these tag clusters (198 bp flanked by 16 bp anchors) in each of four plasmid configurations were designed, constructed, and delivered to an *INS*-1 832/13 rat insulinoma cell line as in Tovar et al. (2023).

### MPRA library analysis

We quantified fragment activity and estimated promoter and position bias using the R package MPRAnalyze, as described in Varshney et al. (2021) and Tovar et al. (2023).

### Elastic net regression

We used elastic net regression to predict MPRA promoter and positional bias from fragment enrichment for a set of 540 nonredundant TF motifs. MPRA bias was represented with Wald-testing-derived z-scores. The set of TF motifs is described in Varshney et al. (2021), and the enrichment quantification was performed as in Tovar et al. (2023). Elastic net regression was performed with the R package “glmnet” (26, 27). We ran the regressions with alpha = 0.8 and alpha = 0 for the position and promoter models, respectively, and 10-fold cross validation.

### Generalized linear models

We modeled total fragment activity across all plasmid promoter and position configurations from strand-of-origin and functional annotations using the R package. Tissue-specific functional annotations were produced using ChromHMM(28).

## RESULTS

### MPRA with position- and promoter-variable configurations

We sought to interrogate position- and promoter-dependent regulatory activity of islet-derived TSSs. We obtained a set of fragments derived from cap analysis of gene expression (CAGE) in human pancreatic islets (29). CAGE works by sequencing 5’ cap-captured mRNAs from the 5’ end, and thus provides a robust method of identifying TSSs (30). The final outputs of CAGEseq processing are CAGE tag clusters, which are distinct groups of overlapping TSSs (31). We defined our fragments as the 198bp forward-strand sequence centered on CAGE tag clusters (Figure 2A). Our final library consisted of 1,305 fragments.

**Figure 2.**
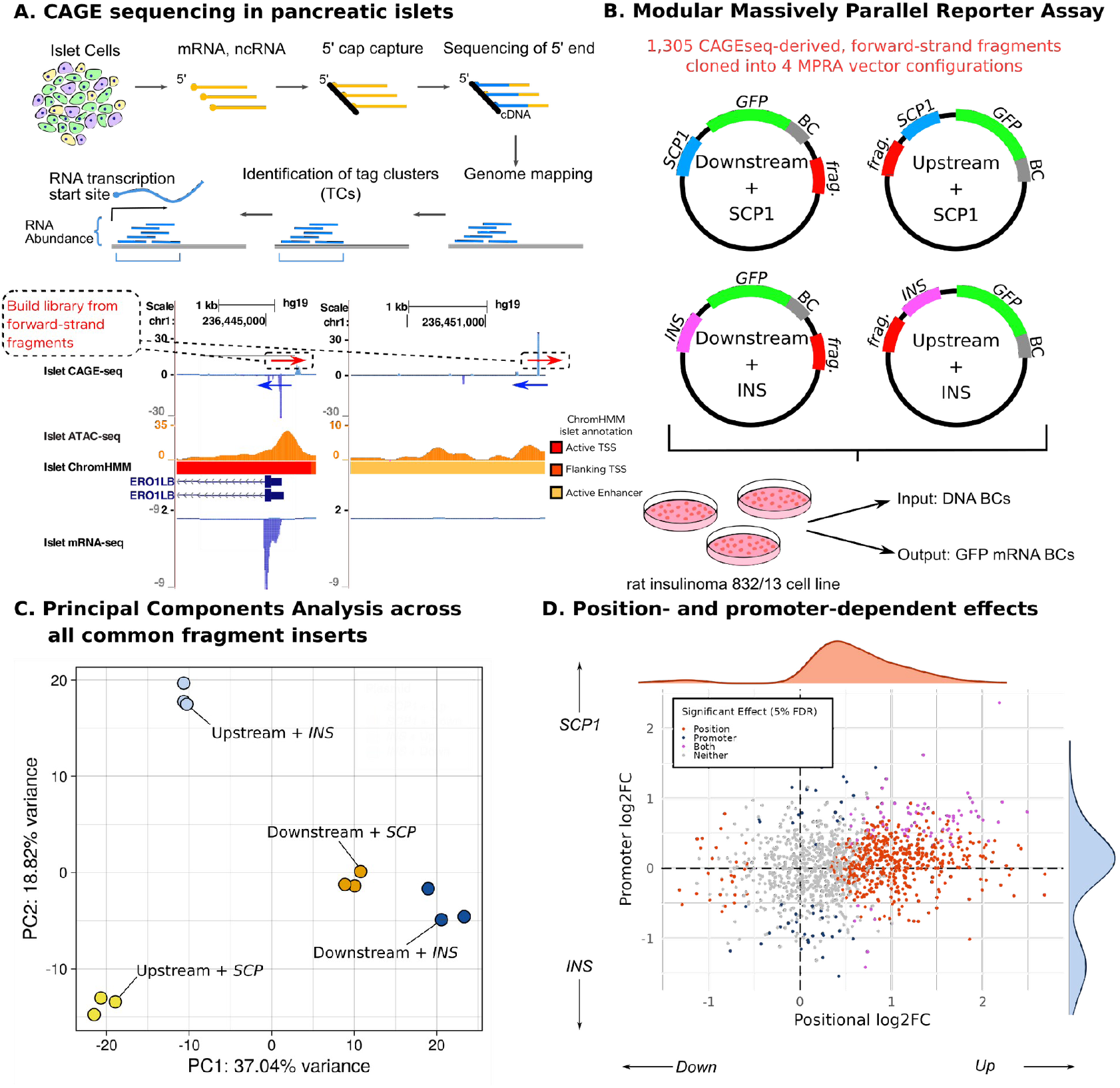
MPRA library construction. (A) Summary of islet CAGE-seq workflow at an example TSS. The genome browser shot shows two CAGE-seq peaks and their associated mRNA-seq, ATAC-seq, and genomic annotation overlaps, all in pancreatic islets. (B) Each fragment was cloned into 4 plasmid configurations: [insert the four descriptions here]. The plasmid library was transfected into the 832/13 rat insulinoma cell line in triplicate. Amplicon sequencing libraries were generated from plasmid DNA (input) and cellular RNA (output). (C) PCA of fragment quantification, where positional configuration is captured primarily by PC1 and promoter is captured by PC2. (D) Distribution of log2FC of enhancer activity between promoter and position conditions for all fragments. Fragments with significant difference in enhancer activity (5% FDR) across the position, promoter, or both conditions are in orange, blue, and purple, respectively.

We cloned each fragment into four different plasmid constructs. Each of the four plasmid configurations utilized one of two potential promoters–either the SCP1 super core promoter or the *INS* human insulin gene promoter–and one of two potential positions–either downstream or upstream of the promoter element (Figure 2B). Each plasmid also received a GFP signal and a unique barcode, which could be used to map back to the fragment and configuration of that plasmid. The plasmid library was transfected into a 832/13 rat insulinoma cell line in triplicate via electroporation. RNA was harvested from these cells 24 hours later and sequenced along with the input plasmid DNA.

We used the R package MPRAnalyze to quantify the activity of each fragment as the ratio of the amount of barcode RNA to the amount of barcode DNA (32). Principal component analysis of fragment activity revealed that, as expected, fragments clustered together based on their plasmid configuration. The first principal component further separated clusters based on position (37.04% of variance). The upstream clusters then separated across principal component 2 (18.82% of variance) by promoter (Figure 2C). Analyzing the fold-change of activity between the different configuration contrasts revealed 565 fragments with significant positional bias, and 124 fragments with significant promoter bias (Figure 2D). Notably, a majority (95%, n = 539/565) of all fragments with significant positional effects (5% FDR) were biased towards having higher activity when placed upstream of the promoter element. 7

### MPRA demonstrates promoter- and position-dependent transcription factor binding site motifs

Given the observed position- and promoter-bias in our fragment library, we sought to identify the sequence motifs driving these biases. For each fragment, we used a Wald test to create a position bias score–where upstream-biased fragments have a more positive score and downstream-biased fragments have a more negative score–and a promoter bias score–where SCP1-biased fragments have a more positive score and *INS*-biased fragments have a more negative score. We performed an elastic net regression with 10-fold cross-validation to predict these bias scores from fragment overlap with a condensed set of 420 non-redundant transcription factor binding site (TFBS) motifs (33) (Figure 3A). Given the large input feature set, elastic net regression is ideal for this analysis, as it can shrink the this set to be more manageable and informative without removing too many important features (34) (Complete model coefficient lists for the position and promoter models are in Supplementary Data 2 and 3, respectively).

**Figure 3.**
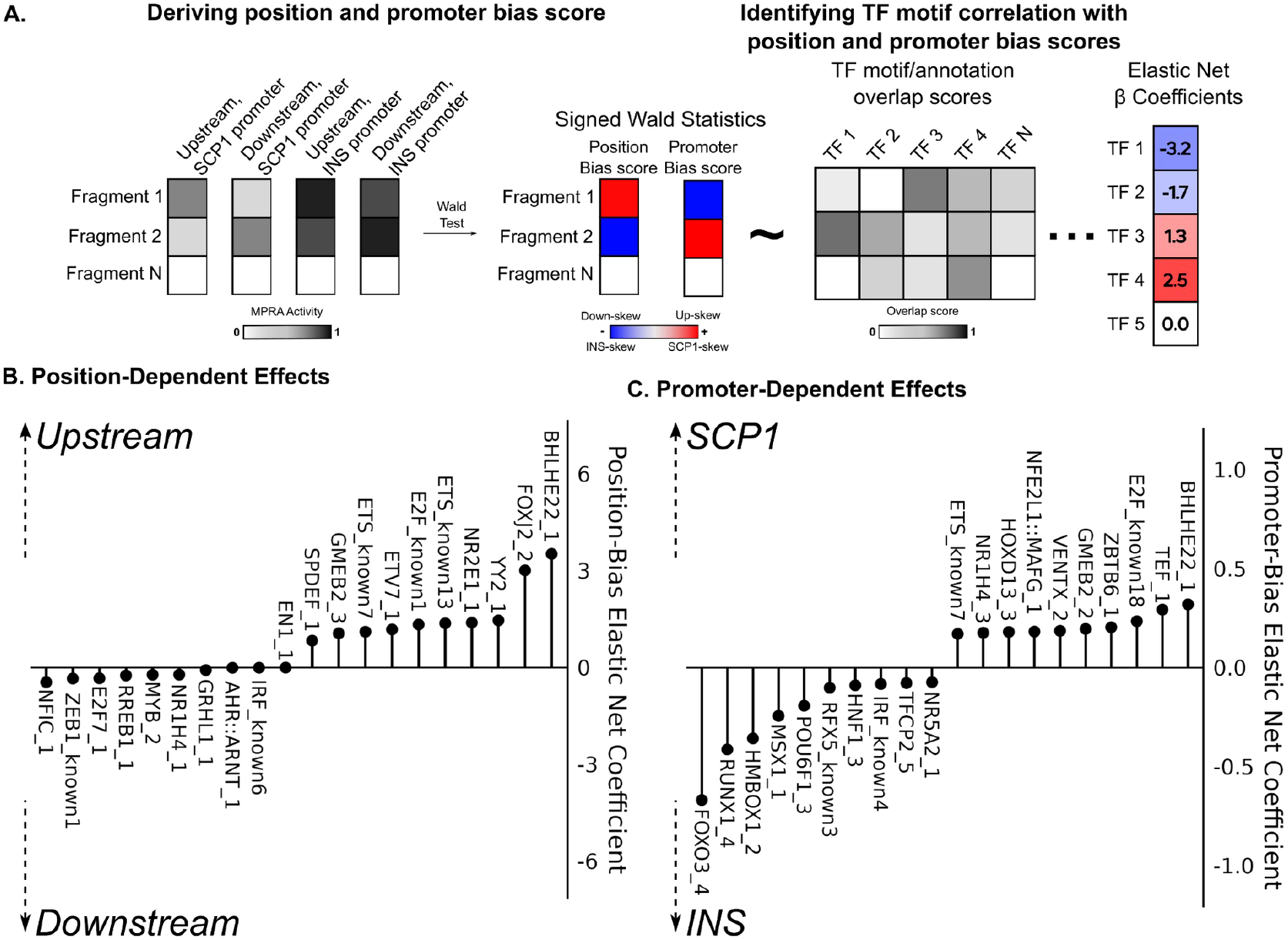
(A) We jointly modeled fragment-level activity with position and promoter as coefficients using MPRAnalyze. To estimate effects of each coefficient, we performed a Wald test, then used signed Wald statistics as our position and promoter bias scores. An elastic net model was used to predict per-fragment position and promoter bias scores on fragment from overlap with TF motif scores. (B) 10 most positive and most negative motif coefficients for position elastic net model. A more positive score indicates that the motif biases fragments to have higher activity when the fragment is upstream of the promoter. A more negative score indicates that the motif biases fragments to have higher activity when the fragment is downstream of the promoter (C) 10 most positive and most negative motif coefficients for promoter elastic net model. A more positive score indicates that the motif biases fragments to have higher activity when the fragment is paired with the SCP1 promoter. A more negative score indicates that the motif biases fragments to have higher activity when the fragment is paired with the INS promoter.

The model coefficients for position bias scores, and especially for upstream-biased fragments, were much stronger than the coefficients for promoter bias scores. For example, the presence of BHLHE22 and E2F1 motifs was associated with higher activity in the upstream position and with the SCP1 promoter (Figures 3B, 3C). The overall fragment bias towards higher activity when placed upstream of the promoter element thus seems to recapitulate the original environment of the fragments as transcription start sites, which likewise initiate transcription upstream of their target gene. Meanwhile, fragments containing motifs corresponding to TFs such as NFIC and ZEB1, which are known to bind enhancers, were biased to have higher activity when placed downstream of the promoter and reporter gene (35–37). Additionally, fragments containing the TFBS motif for HNF1A and RFX5 had higher activity when placed alongside the *INS* promoter (Figure 3C). Pathogenic coding variants in HNF1A cause maturity onset diabetes of the young (MODY) (38). T2D GWAS loci are enriched in RFX-family footprints, with RFX6 being predicted to bind in an islet-specific manner (39). That fragments overlapping these TFBS motifs exhibited activity bias when paired with the *INS* promoter is thus likely a recapitulation of their function in beta cells. Taken together, these results clearly demonstrate how MPRA configuration can differentially affect the enhancer activity of different fragments in a sequence-dependent manner, and how these effects can be understood in the native context of the sequences being assayed.

### MPRA demonstrates tissue- and strand-specificity of TSSs

For our library, following standard MPRA protocols, the forward strand sequence was taken for all CAGE-seq tag clusters assayed, regardless of strand-of-origin. We sought to identify to what extent this standard practice affects MPRA readout and thus interpretation of fragment activity. Given that promoters are primarily unidirectional, we expected to see that CAGE-strand concordant fragments would have higher activity than CAGE-strand discordant fragments, regardless of plasmid configuration (40). Additionally, given that the CAGE peaks were identified in pancreatic islets, we expected that overlap with islet chromHMM regulatory element annotations would likewise predict fragment readout.

Our fragments were approximately evenly distributed between having a forward or reverse strand-of-origin, with slightly more (56%, 734/1305) fragments coming from the reverse strand (Figure 4A). These reverse strand fragments were consequently tested with a discordant strandedness in the MPRA. As expected–given that our fragments were constructed from TSSs identified from CAGE sequencing–a majority of our tag clusters overlapped chromHMM-annotated promoters, in addition to islet ATACseq peaks, with a small number (n = 35/1305) overlapping islet enhancers (Figure 4A). We noticed that, for fragments overlapping both ChromHMM annotated promoters and enhancers, fragments that were strand-concordant with their original CAGE tag clusters tended to have higher MPRA activity than fragments that were strand-discordant with their CAGE tag clusters (Figure 4B). We sought to test this more rigorously with generalized linear models (GLMs). Following the background of our data, overlap with islet promoter sites (t = 6.26, p < 0.001) and islet ATACseq-peaks (t = 6.40, p < 0.001) predicted significantly higher fragment activity, while overlap with islet enhancers predicted significantly lower activity (t = -2.60, p < 0.01). Meanwhile, forward strand-of-origin predicted significantly higher MPRA activity (t = 7.34, p < 0.001) (Figure 4C), suggesting that testing elements in their original strand configuration influences measured enhancer activity in an MPRA construct.

**Figure 4.**
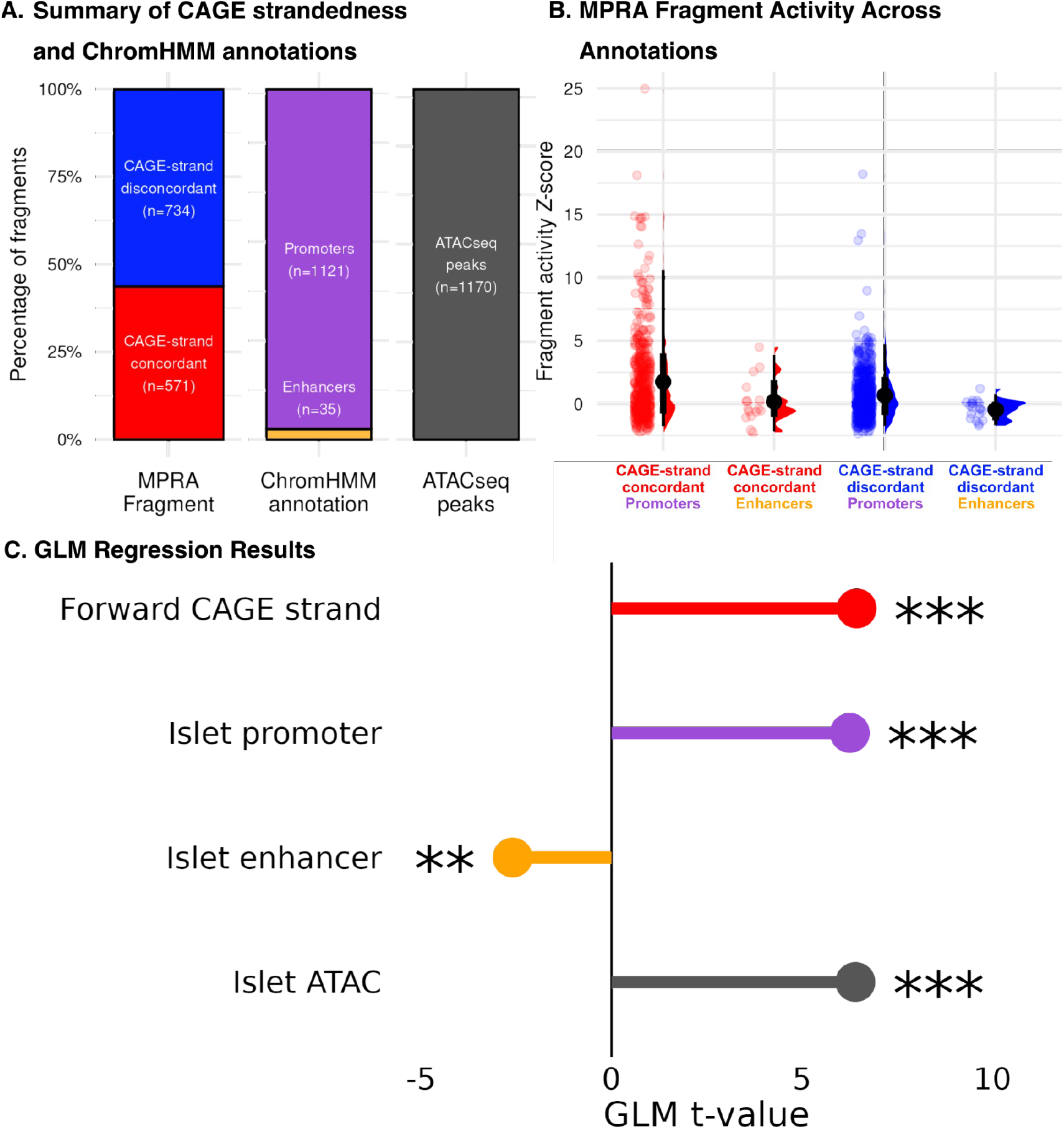
(A) Our fragment library consists of 570 CAGE-strand concordant (forward strand) and 725 reverse CAGE-strand discordant (reverse strand) fragments. Additionally, 1121 fragments overlapped with islet promoters, 35 with islet enhancers, and 1170 with islet ATACseq peaks (B) MPRA activity as a Z-score for fragments split by their CAGE-strand concordance and pancreatic islet ChromHMM annotation category (C) GLM reveals the significant effects of strand-of-origin, in addition to pancreatic islet ChromHMM annotations, on fragment activity (** = p < 0.01, *** = p < 0.001).

## DISCUSSION

We generated an MPRA library from 1,305 CAGE-identified islet TSSs, testing each fragment in 4 different plasmid configurations. These configurations tested fragments alongside two different promoters (the ubiquitously expressed SCP1 promoter and the islet-specific human insulin promoter *INS*), in two different configurations (placed either upstream or downstream of the promoter). We noticed significant promoter- and position-effects, indicating that our islet TSS-derived fragments can exhibit a range of expression activity depending on the configuration of the MPRA plasmid they are tested in. Using elastic net regression, we found strong positional effects for several TFBS motifs. BHLHE22 and FOXJ2 motifs were associated with enhanced expression when placed upstream of the promoter element, while HNF1A and RFX motifs were associated with enhanced expression in the context of the *INS* promoter. We also identified a strong directional effect, with fragments originating from the forward strand demonstrating much higher MPRA activity than fragments originating from the reverse strand.

This dataset has several limitations. The plasmid library was only tested in the rat insulinoma 823/13 cell line, as a model cell line pancreatic beta cells, which was the most tractable option available at the time of experimentation. While this line replicates many features of the transcriptional environment in human pancreatic beta cells (41), future studies will be improved with use of a human-derived beta cell line such as EndoC-βH3 (42). Additionally, a known limitation of plasmid MPRAs is that they lose the “in-genome” effects of chromatin structure (43). The use of an integrated system (e.g., lentiMPRA) in the future could help to overcome this limitation; however, these systems still cannot place sequences in their exact genomic context in a high-throughput manner, making the chromatin context that sequences are placed into somewhat arbitrary. Finally, we have only shown these results for islet transcription start sites and T2D GWAS fragments; these configurational differences may appear less distinct for other fragment libraries (24).

Additionally, at least in the case of forward-strand TSSs, respect to strand-of-origin has a massive effect on fragment activity. Fragments tested in the reverse of their native orientation had much lower activity than fragments tested in their original orientation. This makes it difficult to understand the full range of regulatory activity driven by a given fragment when only tested against a single minimal promoter or in a single plasmid positional configuration, and always in the forward orientation, as is the case in a majority of MPRA experiments. These aspects limit our ability to understand the activity of these fragment sequences in their native genomic and biological context. This result also suggests that additional studies which interrogate the joint influences of cell context and strandedness would enhance our broader understanding of gene regulation.

While these results point to a key limitation of MPRAs, they also demonstrate a solution and path forward. The activity of a given fragment in an MPRA experiment can be significantly altered by the choice of promoter and plasmid positional configuration. This bias is apparently sequence-specific, being driven in one direction or the other by specific TF motifs. Future MPRA experiments should carefully consider the activity of fragments across multiple plasmid configurations. These configurations should be informed by the biological contexts in which their fragments of interest are relevant. With this strategy, researchers may be able to produce more informative, biologically meaningful results.

## DATA AVAILABILITY

The raw and processed data from this study is available at the GEO accession GSE279057. The code used to preprocess and analyze data is adapted from a previous publication (Varshney et al. 2021) and is found at https://github.com/ParkerLab/STARR-seq-Analysis-Pipeline. The code used to run the elastic net and generalized linear models is found at https://github.com/boseHere/TSS_MPRA_2025.

## ACKNOWLEDGEMENTS

The authors of this paper would like to acknowledge the support and feedback provided by members of the Parker and Kitzman labs (University of Michigan). Additionally, we thank the University of Michigan Advanced Genomics Core for providing sequencing services.

