## Supplemental Data 1 for "Massively parallel reporter assay reveals promoter-, position-, and strand-specific effects in transcription start sites"

Supplementary Data 1: Review of 146 published MPRA studies, annotated by the number of different promoter elements and plasmid configurations each fragment was tested with. List of studies was generated using the Pubmed search (("massively parallel reporter assay") NOT (Review[Publication Type]) NOT (Meta-Analysis[Publication Type])) on Aug. 29, 2025. Databases, protocols, meta-analyses, and analyses that do not create new data were removed.

| PMID | DOI | Number of promoters tested alongside fragment | Number of positions (plasmid configurations) fragment was tested in |
|---|---|---|---|
| 35298243 | 10.1126/science.abj5117 | 1 | 1 |
| 37413987 | 10.1016/j.cell.2023.06.007 | 1 | 1 |
| 34534445 | 10.1016/j.cell.2021.08.025 | 1 | 1 |
| 30033119 | 10.1016/j.stem.2018.06.014 | 3 | 1 |
| 37071996 | 10.1016/j.devcel.2023.03.020 | 1 | 1 |
| 27259153 | 10.1016/j.cell.2016.04.027 | 1 | 1 |
| 36705030 | 10.1161/CIRCULATIONAHA.122.061955 | 1 | 1 |
| 34650237 | 10.1038/s41588-021-00947-3 | 1 | 1 |
| 37387536 | 10.1111/tpj.16373 | 1 | 1 |
| 37953348 | 10.1038/s42003-023-05483-w | 1 | 1 |
| 36156153 | 10.1093/nar/gkac806 | 1 | 1 |
| 31530582 | 10.1101/gr.247312.118 | 1 | 2 |
| 28204611 | 10.1093/nar/gkw942 | 1 | 1 |
| 37996647 | 10.1038/s41556-023-01296-5 | 1 | 1 |
| 30158147 | 10.1101/gr.231886.117 | 2 | 1 |
| 36646877 | 10.1038/s41593-022-01243-x | 1 | 1 |
| 38407202 | 10.7554/eLife.89371 | 1 | 1 |
| 34849835 | 10.1093/g3journal/jkab404 | 1 | 1 |
| 33885362 | 10.7554/eLife.63713 | 1 | 1 |
| 37658059 | 10.1038/s41467-023-41081-4 | 1 | 1 |
| 29225039 | 10.1016/j.molcel.2017.11.014 | 1 | 1 |
| 38389303 | 10.1016/j.xhgg.2024.100279 | 1 | 1 |
| 31152051 | 10.1101/gr.242552.118 | 1 | 1 |
| 30451991 | 10.1038/nbt.4285 | 2 | 1 |
| 29410437 | 10.1038/s41467-018-02980-z | 1 | 2 |
| 36777181 | 10.1016/j.xgen.2022.100234 | 1 | 2 |
| 37087538 | 10.1038/s41467-023-37960-5 | 1 | 1 |
| 29889606 | 10.1080/21541264.2018.1486150 | 1 | 1 |
| 36107770 | 10.1093/nar/gkac763 | 1 | 1 |
| 35534523 | 10.1038/s41598-022-11589-8 | 1 | 1 |
| 33357440 | 10.1016/j.celrep.2020.108531 | 1 | 1 |
| 38183988 | 10.1016/j.ajhg.2023.12.008 | 1 | 1 |
| 36763080 | 10.7554/eLife.71235 | 2 | 1 |
| 38365907 | 10.1038/s41598-024-54302-7 | 1 | 1 |
| 36192170 | 10.1101/gr.276863.122 | 1 | 1 |
| 28525990 | 10.1186/s12864-017-3785-4 | 1 | 1 |

| | | | |
|---|---|---|---|
| 34978147 | 10.1002/alz.12534 | 1 | 1 |
| 36555130 | 10.3390/ijms232415485 | 1 | 1 |
| 37868037 | 10.1016/j.xgen.2023.100404 | 1 | 1 |
| 33712590 | 10.1038/s41467-021-21854-5 | 1 | 1 |
| 33970899 | 10.1371/journal.pcbi.1008982 | 1 | 1 |
| 26713262 | 10.7717/peerj.1527 | 1 | 1 |
| 37492106 | 10.1016/j.xgen.2023.100330 | 1 | 1 |
| 39818206 | 10.1016/j.devcel.2024.12.038 | 1 | 1 |
| 35866592 | 10.1093/gbe/evac108 | 1 | 1 |
| 32103011 | 10.1038/s41467-020-14853-5 | 1 | 1 |
| 22371084 | 10.1038/nbt.2137 | 1 | 1 |
| 31631012 | 10.1016/j.stem.2019.09.010 | 1 | 1 |
| 38647082 | 10.1093/nar/gkae285 | 1 | 1 |
| 22371081 | 10.1038/nbt.2136 | 1 | 1 |
| 33849996 | 10.2337/db20-1087 | 1 | 1 |
| 32239644 | 10.15252/emmm.202012112 | 1 | 1 |
| 34489471 | 10.1038/s41467-021-25614-3 | 1 | 1 |
| 32426415 | 10.1016/j.omtm.2020.04.006 | 1 | 1 |
| 34662402 | 10.1093/molbev/msab304 | 1 | 1 |
| 33626337 | 10.1016/j.ajhg.2021.02.006 | 1 | 1 |
| 23512712 | 10.1101/gr.144899.112 | 1 | 1 |
| 34850108 | 10.1093/nar/gkab1100 | 1 | 1 |
| 32133495 | 10.1093/nar/gkaa147 | 1 | 1 |
| 37516102 | 10.1016/j.celrep.2023.112840 | 1 | 1 |
| 31164647 | 10.1038/s41467-019-10439-y | 1 | 1 |
| 35082832 | 10.3389/fgene.2021.785934 | 1 | 1 |
| 31464371 | 10.15252/msb.20198875 | 1 | 1 |
| 34390653 | 10.1016/j.ajhg.2021.07.009 | 1 | 1 |
| 29728462 | 10.1073/pnas.1722055115 | 7 | 1 |
| 31503409 | 10.1002/ajmg.b.32761 | 1 | 1 |
| 30537984 | 10.1186/s13059-018-1589-8 | 1 | 1 |
| 27831498 | 10.1101/gr.212092.116 | 1 | 1 |
| 37879864 | 10.1261/rna.079752.123 | 1 | 1 |
| 38442181 | 0.1073/pnas.2309469121 | 1 | 1 |
| 33179598 | 10.7554/eLife.62669 | 1 | 1 |
| 36947129 | 10.7554/eLife.83593 | 1 | 1 |
| 37906604 | 10.1371/journal.pgen.1011014 | 1 | 1 |
| 27259154 | 10.1016/j.cell.2016.04.048 | 1 | 1 |
| 34663436 | 10.1186/s13059-021-02509-6 | 1 | 1 |
| 38997252 | 10.1038/s41467-024-50174-7 | 1 | 1 |
| 40393459 | 10.1016/j.xgen.2025.100882 | 1 | 1 |
| 27078102 | 10.1073/pnas.1602886113 | 2 | 1 |
| 31344026 | 10.1371/journal.pgen.1008287 | 1 | 1 |
| 31227602 | 10.1101/gr.245159.118 | 1 | 1 |
| 25340400 | 10.1371/journal.pgen.1004592 | 1 | 1 |
| 27783940 | 10.1016/j.celrep.2016.09.066 | 2 | 1 |
| 38177677 | 10.1038/s41594-023-01171-9 | 1 | 1 |
| 32043966 | 10.7554/eLife.41279 | 1 | 1 |
| 33046894 | 10.1038/s41592-020-0965-y | 1 | 2 |
| 34475398 | 10.1038/s41467-021-25514-6 | 1 | 1 |

| | | | |
|---|---|---|---|
| 35315433 | 10.1038/s41467-022-28659-0 | 1 | 1 |
| 36834916 | 10.3390/ijms24043509 | 1 | 1 |
| 23328393 | 10.1126/science.1232542 | 1 | 1 |
| 23892608 | 10.1038/ng.2713 | 1 | 1 |
| 25872643 | 10.1038/ncomms7905 | 1 | 1 |
| 26486725 | 10.1101/gr.191593.115 | 1 | 1 |
| 26576614 | 10.1101/gr.193789.115 | 1 | 1 |
| 27311442 | 10.1101/gr.204834.116 | 1 | 2 |
| 27524623 | 10.1016/j.celrep.2016.07.050 | 1 | 1 |
| 27565349 | 10.1016/j.cell.2016.07.049 | 1 | 1 |
| 27667684 | 10.1016/j.cell.2016.08.071 | 1 | 1 |
| 27701403 | 10.1038/nbt.3678 | 1 | 1 |
| 28137873 | 10.1073/pnas.1621150114 | 1 | 1 |
| 28973438 | 10.1093/nar/gkx577 | 1 | 1 |
| 29061142 | 10.1186/s13059-017-1322-z | 1 | 1 |
| 29151363 | 10.1186/s13059-017-1345-5 | 1 | 1 |
| 29256496 | 10.1038/nmeth.4534 | 2 | 1 |
| 29789573 | 10.1038/s41467-018-04451-x | 1 | 1 |
| 30045748 | 10.1186/s13059-018-1473-6 | 1 | 1 |
| 30568279 | 10.1038/s41467-018-07746-1 | 1 | 1 |
| 31267113 | 10.1038/s41587-019-0164-5 | 1 | 1 |
| 31395865 | 10.1038/s41467-019-11526-w | 1 | 1 |
| 32248749 | 10.1161/ CIRCRESAHA.119.316006 | 1 | 1 |
| 32483191 | 10.1038/s41467-020-16590-1 | 1 | 1 |
| 32616518 | 10.1101/gr.260463.119 | 1 | 1 |
| 32747698 | 10.1038/s41398-020-00953-9 | 1 | 1 |
| 39848247 | 10.1016/j.cell.2024.12.022 | 1 | 1 |
| 39443793 | 10.1038/s41586-024-08070-z | 1 | 1 |
| 38378865 | 10.1038/s41588-024-01669-y | 1 | 1 |
| 39317738 | 10.1038/s41588-024-01896-3 | 1 | 1 |
| 39644900 | 10.1016/j.cels.2024.11.003 | 3 | 1 |
| 38724816 | 10.1007/s10517-024-06074-3 | 1 | |
| 38947339 | 10.3389/fimmu.2024.1387253 | 1 | 1 |
| 39631147 | 10.1016/j.ebiom.2024.105480 | 1 | 1 |
| 38773080 | 10.1038/s41467-024-48436-5 | 1 | 1 |
| 39609378 | 10.1038/s41467-024-54502-9 | 3 | 1 |
| 39738051 | 10.1038/s41467-024-55274-y | 1 | 1 |
| 40205616 | 10.1186/s13073-025-01459-z | 1 | 1 |
| 40846081 | 10.1016/j.jaci.2025.07.032 | 1 | 1 |
| 38413607 | 10.1038/s41531-024-00659-5 | 1 | 1 |
| 40680142 | 10.1126/sciadv.ads9164 | 1 | 1 |
| 38334359 | 10.7554/eLife.85235 | 1 | 1 |
| 39532105 | 10.1016/j.devcel.2024.10.017 | 1 | 1 |
| 40670354 | 10.1038/s41467-025-61734-w | 1 | 1 |
| 40494627 | 10.1101/gr.280320.124 | 1 | 1 |
| 39995040 | 10.1093/nar/gkaf097 | 1 | 1 |
| 40414878 | 10.1186/s13059-025-03610-w | 2 | 1 |
| 40715118 | 10.1038/s41467-025-62000-9 | 1 | 1 |
| 40738258 | 10.1016/j.jgg.2025.07.008 | 1 | 1 |

| | | | |
|---|---|---|---|
| 40838804 | 10.1093/g3journal/jkaf192 | 1 | 1 |
| 39737967 | 10.1038/s41467-024-54723-y | 1 | 1 |
| 40586305 | 10.1093/nar/gkaf568 | 2 | 1 |
| 40399339 | 10.1038/s41467-025-60023-w | 1 | 1 |
| 40318978 | 10.1016/j.ard.2025.04.001 | 1 | 1 |
| 39964837 | 10.7554/eLife.97682 | 1 | 1 |
| 40485594 | 10.1093/plcell/koaf084 | 1 | 1 |
| 40210244 | 10.1093/nar/gkaf224 | 1 | 1 |
| 40659498 | 10.1101/gr.279957.124 | 1 | 1 |
| 40274775 | 10.1038/s41467-025-58970-5 | 1 | 1 |
| 38997781 | 10.1186/s12920-024-01954-z | 1 | 1 |
