## Supplemental Data 3 for "Massively parallel reporter assay reveals promoter-, position-, and strand-specific effects in transcription start sites"

Supplementary Data 3: Coefficients from elastic net model predicting promoter bias from overlap with transcription-factor binding site motifs. A more positive coefficient indicates that fragments with the motif have higher plasmid activity with the SCP promoter, while a more negative coefficient indicates that fragments with the motif have higher plasmid activity with the *INS* promoter.

| **Model coefficient** | **Transcription Factor Binding Site Motif** |
|---|---|
| 0.319964616859408 | BHLHE22_1 |
| 0.293524388458332 | TEF_1 |
| 0.234710461985691 | E2F_known18 |
| 0.204263463417694 | ZBTB6_1 |
| 0.19603984934424 | GMEB2_2 |
| 0.186138890545583 | VENTX_2 |
| 0.182271895151008 | `NFE2L1::MAFG_1` |
| 0.180608817004959 | HOXD13_3 |
| 0.176309707596735 | NR1H4_3 |
| 0.170745518047966 | ETS_known7 |
| 0.157091898449797 | FOXJ2_2 |
| 0.149301928977088 | PBX1_1 |
| 0.137504780183651 | E2F_known1 |
| 0.129331877976445 | DMRT1_1 |
| 0.122881725560097 | GMEB2_3 |
| 0.110419370453192 | `NKX2-5_3` |
| 0.0908973985377944 | EGR1_known9 |
| 0.0890163322533468 | ETS_known13 |
| 0.0882374313481226 | YY1_known1 |
| 0.0795614493185918 | TCF7L2_known1 |
| 0.0762473460268454 | `DDIT3::CEBPA_1` |
| 0.0744338700514282 | HINFP_1 |
| 0.0653258211672438 | J4153 |
| 0.0652873796625112 | FEV_1 |
| 0.0630307763011365 | ZEB1_known4 |
| 0.0542448861045126 | ETV7_1 |
| 0.0537007683406604 | SOX17_2 |
| 0.0502524385823001 | NRF1_known2 |
| 0.0428308477687616 | YY2_1 |
| 0.0427824552663358 | CPHX_1 |
| 0.038493459562817 | HIC1_5 |
| 0.0365303915255531 | CUX1_6 |
| 0.0330455322662289 | YY1_known4 |
| 0.0326563585595943 | IRF_known16 |
| 0.0325499881255523 | TBX5_5 |
| 0.0314920344104799 | ZNF219_1 |
| 0.026489420955191 | FLI1_4 |
| 0.026085080337118 | ITGB2_1 |

| | |
|---|---|
| 0.024932372411848 | ZNF143_known2 |
| 0.0234500185697827 | TBX21_6 |
| 0.0233745362309738 | PAX2_1 |
| 0.0232530391387783 | KLF13_1 |
| 0.0207734236000009 | HNF4_known26 |
| 0.0194351370098085 | PPARA_1 |
| 0.0138849535038736 | E2F_known2 |
| 0.0137622162484968 | ZNF8_1 |
| 0.012743882832353 | RXRG_3 |
| 0.011653148498354 | AHR_2 |
| 0.0102925479948754 | SOX1_1 |
| 0.00687872900643944 | SMAD4_1 |
| 0.00664429233093375 | ETS_known1 |
| 0.00530858741688046 | E2F_known23 |
| 0.000210671864537499 | E2F_known15 |
| -0.00426912707676313 | MYBL2_1 |
| -0.00652962562299895 | HSFY2_2 |
| -0.0118848056021666 | RREB1_2 |
| -0.0242123397554972 | GTF2I_1 |
| -0.0259329693880413 | SREBP_known4 |
| -0.0278121794986198 | MEF2_known1 |
| -0.0516010887401002 | SRY_6 |
| -0.05519622820781 | MZF1_2 |
| -0.0566318611088047 | SREBP_known2 |
| -0.0703110172293828 | INSM1_1 |
| -0.0732129653287389 | NR5A2_1 |
| -0.0773233738465818 | TFCP2_5 |
| -0.0821503790409992 | IRF_known4 |
| -0.0887572281668028 | HNF1_3 |
| -0.100996617243515 | RFX5_known3 |
| -0.191864837828888 | POU6F1_3 |
| -0.243220201327591 | MSX1_1 |
| -0.35691978758685 | HMBOX1_2 |
| -0.412821128735226 | RUNX1_4 |
| -0.668489908611629 | FOXO3_4 |
