## Supplemental Data 2 for "Massively parallel reporter assay reveals promoter-, position-, and strand-specific effects in transcription start sites"

Supplementary Data 2: Coefficients from elastic net model predicting position bias from overlap with transcription-factor binding site (TFBS) motifs. A more positive coefficient indicates that fragments with the motif have higher plasmid activity in the upstream configuration, while a more negative coefficient indicates that fragments with the motif have higher plasmid activity in the downstream configuration.

| **Model coefficient** | **Transcription Factor Binding Site Motif** |
|---|---|
| 3.52562996176848 | BHLHE22_1 |
| 3.01497004058349 | FOXJ2_2 |
| 1.46373927636467 | YY2_1 |
| 1.39376099466038 | NR2E1_1 |
| 1.37499562178328 | ETS_known13 |
| 1.33562589031284 | E2F_known1 |
| 1.18321296532986 | ETV7_1 |
| 1.111182767387 | ETS_known7 |
| 1.0603185394601 | GMEB2_3 |
| 0.834796942353679 | SPDEF_1 |
| 0.79638668598655 | ZNF143_known2 |
| 0.749971766187846 | `NKX2-5_3` |
| 0.72463559360545 | VENTX_2 |
| 0.680933478499298 | NRF1_known2 |
| 0.658746625595083 | ZNF8_1 |
| 0.618219196152637 | E2F_known18 |
| 0.614670634596061 | YY1_known4 |
| 0.610346064006302 | FEV_1 |
| 0.605373454050388 | T_2 |
| 0.560754949733233 | HINFP_1 |
| 0.536288449402198 | RORA_2 |
| 0.522227354525412 | E2F_known2 |
| 0.496979582480918 | FOXB1_1 |
| 0.469412507538722 | YY1_known1 |
| 0.383538162664215 | E2F_known23 |
| 0.337682023833318 | NR3C1_known13 |
| 0.318763350664279 | GMEB2_2 |
| 0.310562017318377 | RARG_8 |
| 0.274213277970278 | GLI2_2 |
| 0.227435738335378 | IRF_known2 |
| 0.217991874333012 | ETV6_1 |
| 0.211273996625631 | ARNTL_1 |
| 0.208619890638352 | HIF1A_1 |
| 0.187387875012351 | OSR2_1 |
| 0.151413925419386 | SMAD_1 |
| 0.151064668557514 | IRF_known10 |
| 0.146908595490574 | EGR1_known9 |

| | |
|---|---|
| 0.130150171843634 | CUX1_6 |
| 0.113668922939779 | ATF3_known5 |
| 0.0978633071316993 | NR1H4_3 |
| 0.0753074223720239 | SP4_2 |
| 0.0670299999647747 | GFI1_1 |
| 0.0649585846780402 | SPI1_known1 |
| 0.0417198905083158 | RORA_7 |
| 0.0406914524636692 | IRF_known16 |
| 0.0328258673258894 | KLF13_1 |
| 0.0140713940816713 | HNF4_known26 |
| 0.000191995292464116 | EN1_1 |
| -0.00345412472045617 | IRF_known6 |
| -0.00530653286737288 | `AHR::ARNT_1` |
| -0.0763846986188357 | GRHL1_1 |
| -0.218241139180648 | NR1H4_1 |
| -0.220717965405663 | MYB_2 |
| -0.244008651051658 | RREB1_1 |
| -0.330204290935185 | E2F7_1 |
| -0.337958969051004 | ZEB1_known1 |
| -0.451622299464286 | NFIC_1 |
